# New genes and functional innovation in mammals

**DOI:** 10.1101/090860

**Authors:** José Luis Villanueva-Cañas, Jorge Ruiz-Orera, M.Isabel Agea, Maria Gallo, David Andreu, M.Mar Albà

## Abstract

The birth of genes that encode new protein sequences is a major source of evolutionary innovation. However, we still understand relatively little about how these genes come into being and which functions they are selected for. To address these questions we have obtained a large collection of mammalian-specific gene families that lack homologues in other eukaryotic groups. We have combined gene annotations and *de novo* transcript assemblies from 30 different mamalian species, obtaining about 6,000 gene families. In general, the proteins in mammalian-specific gene families tend to be short and depleted in aromatic and negatively charged residues. Proteins which arose early in mammalian evolution include milk and skin polypeptides, immune response components, and proteins involved in reproduction. In contrast, the functions of proteins which have a more recent origin remain largely unknown, despite the fact that these proteins also have extensive proteomics support. We identify several previously described cases of genes originated *de novo* from non-coding genomic regions, supporting the idea that this mechanism frequently underlies the evolution of new protein-coding genes in mammals. Finally, we show that most young mammalian genes are preferentially expressed in testis, suggesting that sexual selection plays an important role in the emergence of new functional genes.

## INTRODUCTION

The genome and mRNA sequencing efforts of the last two decades have resulted in gene catalogues for a large number of species (Genome 10K Community of Scientists. 2009; Lindblad-Toh et al. 2011; Flicek et al. 2014). This has spurred the comparison of genes across species and the identification of a surprisingly large number of proteins that appear to be limited to one species or lineage (Wood et al. 2002; Domazet-Loso and Tautz 2003; Albà and Castresana 2005; Milde et al. 2009; Tautz and Domazet-Lošo 2011; Toll-Riera, Bostick, et al. 2012; Neme and Tautz 2013; Wissler et al. 2013; Arendsee et al. 2014; Palmieri et al. 2014; Schlötterer 2015). Although these proteins probably hold the key to many recent adaptations (Zhang and Long 2014), they remain, for the most part, still uncharacterized (McLysaght and Hurst 2016).

Species- or lineage-specific orphan genes are defined by their lack of homologues in other species. They may arise by rearrangements of already existing coding sequences, often including partially duplicated genes and/or transposable elements (Zhang et al. 2004; Toll-Riera et al. 2009; Toll-Riera et al. 2011), or completely *de novo* from previously non-coding genomic regions (Levine et al. 2006; Cai et al. 2008; Heinen et al. 2009; Knowles and McLysaght 2009; Toll-Riera et al. 2009; Tautz and Domazet-Lošo 2011; Wu et al. 2011; Reinhardt et al. 2013; McLysaght and Hurst 2016). The latter process is facilitated by the high transcriptional turnover of the genome, which continuously produces transcripts that can potentially acquire new functions and become novel protein coding-genes (Zhao et al. 2014; Ruiz-Orera et al. 2015; Neme and Tautz 2016).

Lineage-specific genes are expected to be major drivers of evolutionary innovation. A well-known example is the nematocyst in Cnidaria that is used to inject toxin into the preys. Some of the major constituents of the nematocyst are Cnidaria-specific proteins, indicating that the birth of new genes in a Cnidaria ancestor was intimately linked to the emergence of a novel trait (Milde et al. 2009). Lineage- or species-specific genes also comprise a large proportion of the proteins specific of Molluscs shells (Aguilera et al. 2017). In mammals, diverse adaptations have also been related to new genes. For example, the caseins, which transport calcium in the milk, originated from duplications of calcium-binding phosphoprotein which underwent very drastic sequence changes early in the evolution of the mammalian group (Kawasaki et al. 2011).

In mammals, there have been a number of systematic studies aimed at the identification of recently evolved genes, particulary in human and to a lesser extent mouse (Knowles and McLysaght 2009; Toll-Riera et al. 2009; Wu et al. 2011; Murphy and McLysaght 2012; Neme and Tautz 2013; Guerzoni and McLysaght 2016). These works have provided some of the first examples of genes likely to have originated *de novo*. However, our knowledge of how these genes have impacted mammalian biology has lagged behind. To fill this gap we have generated a comprehensive list of gene families which have originated at different times during the evolution of mammals. The combinaton of gene expression data, functional annotation, proteomics, and amino acid sequence properties provides novel insights into the birth of new genes in mammals.

## RESULTS

### Identification of mammalian-specific gene families

Our first goal was to build a comprehensive catalogue of mammalian-specific protein-coding gene families. To ensure consistency, our analysis was based on 30 mammalian species with a complete genome sequence, gene annotations and RNA sequencing data (RNA-Seq) (Table S1). The RNA-Seq data was used to assemble transcripts *de novo*, which provided a source of expressed sequences for each species that was independent of the gene annotations and which served to complement them.

First, in each of the 30 species, we selected those genes that lacked homologues in non-mammalian species using BLASTP sequence similarity searches against a panel of 34 species including vertebrates, insects, plants, fungi, and bacteria (Table S1). Subsequently, we eliminated any genes that had initially been classified as mammalian-specific, but had indirect homology, through paralogues, in more distant species. This was done to minimize the inclusion of very rapidly evolving duplicated genes (Toll-Riera et al. 2009).

Next, we used the remaining mammalian-specific genes in each species to perform all against all sequence similarity searches. We clustered the genes into families by iteratively inspecting the lists of sequence similarity hits until no more homologues could be added to a given family. Then we assigned each family to the node that connected the most distant species present in the family. This node was the point from which no further ancestors could be traced back.

A method that relies solely on gene annotations is likely to miss homologues in the species which are not very well annotated. For this reason we performed additional searches of all members of a family against the novel transcripts generated with the RNA-Seq data. In most families the use of RNA-Seq data expanded the range of species with evidence of expression of the gene (Figure S1). In some cases it also resulted in a deepening of the node of origin; approximately 22% of the genes initially classified as species-specific were reclassified into multi-species families. To ensure robustness, in each node we only considered the gene families that contained sequences from at least half of the species derived from that node. The procedure resulted in 2,034 multi-species gene families, altogether containing 10,991 different proteins. We also obtained 3,972 species-specific families, containing 4,173 different proteins. The complete catalogue of gene families, protein sequences, and transcript assemblies is provided as supplementary material.

Figure 1 shows the distribution of gene families in the different nodes of the mammalian tree. About one fourth of the families (439) had a basal origin, they had originated before the split of the major mammalian groups, probably more than 100 Million years ago (class ‘mam-basal’ or 2; nodes 1, 2, 4, 5 and 6 in Figure S2). The rest of multi-species families corresponded to more recent nodes and were classified as ‘mam-young (class 1). This included, for example, the 269 families which were specific of the Catarrhini (old World monkeys and apes, node 21 in Figure S2). These genes were present in macaque and/or baboon, and in the great apes, but were absent from other primate or mammalian branches. Other examples of large sets of phylogenetically-restricted gene families corresponded to the primates (135 families, node 7), the muridae (173 families, node 23) or the felids (52 families, node 14).

**Figure 1.**
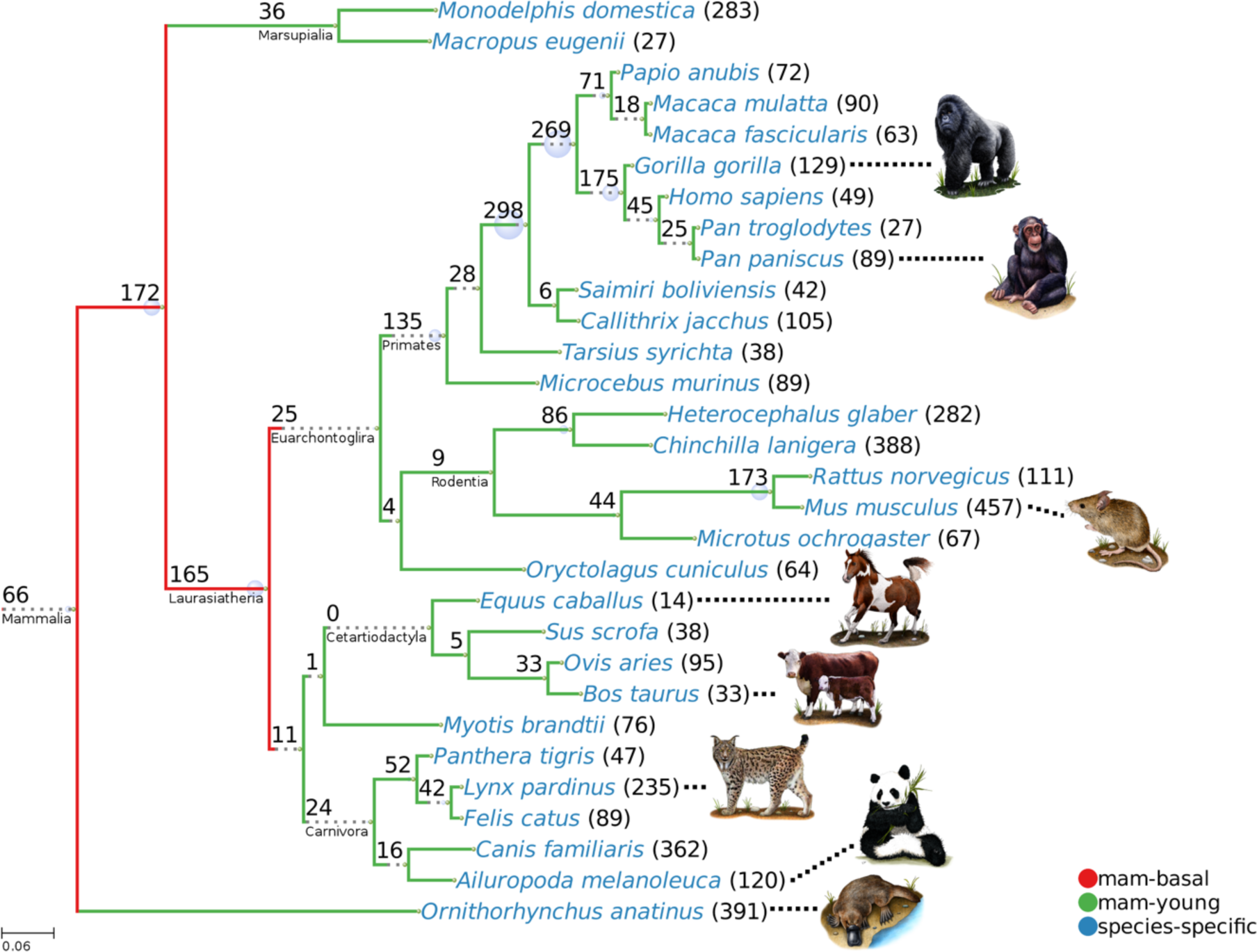
Mammalian tree and number of mammalian-specific gene families. The tree depicts the phylogenetic relationships between 30 mammalian species from different major groups. The values in each node indicate the number of families that were mapped to the branch ending in the node. We define three conservation levels: ‘mam-basal’ (class 2, approximately older than 100 Million years, red), ‘mam-young’ (class 1, green) and ‘species-specific (class 0, blue). The branch length represents the approximate number of substitutions per site as inferred from previous studies (see Methods). The scale bar on the bottom left corner represents 6 substitutions per 100 nucleotides. Dotted lines have been added to some branches to improve readability.

Our dataset included several previously described mammalian-specific genes. One example was neuronatin, an imprinted mammalian-specific gene involved in the regulation of ion channels during brain development (Evans et al. 2005). We found members of this family in 26 placental species but not in marsupials or Monotremata. Another example was the abundant antimicrobial salivary peptide mucin-7 (Bobek and Situ 2003). This gene likley originated in a placental mammal ancestor and evolved under positive selection in response to pathogens (Xu et al. 2016). Another example was dermcidin, a primate-specific antimicrobial peptide secreted in the skin (Toll-Riera et al. 2009). An illustrative list of genes, including these examples and other cases discussed in the next sections, is available from Table 1.

**Table 1.** **Examples of mammalian-specific genes families.** We indicate the node under which we classified the gene. Tree Node numbers: Mammalia 1; Theria 2; Eutheria 4; Primates 7; Haplorrhini 16; Great apes 26; Hum/Chimp 28.

| Gene Name | Description | Tree Node | Features | References |
| --- | --- | --- | --- | --- |
| SCGB | secretoglobin | Mammalia | Gene family, modulation of inflammation | (Jackson et al. 2011) |
| PRM3 | protamine 3 | Theria | Affects sperm motility | (Grzmil et al. 2008) |
| CSN1S1 | casein alpha s1 | Eutheria | Ca-sensitive milk protein, related to vertebrate calcium-binding protein SPARCL1 | (Kawasaki et al. 2011) |
| LCE6A | late cornified envelope 6A | Eutheria | Formation of the skin, part of the epidermal differentiation complex | (Strasser et al. 2014) |
| IL2 | interleukin 2 | Eutheria | Cytokine, rapid sequence divergence | (Bird et al. 2005) |
| MUC7 | mucin 7 | Eutheria | Antimicrobial peptide, secreted in saliva | (Bobek and Situ 2003; Xu et al. 2016) |
| NNAT | neuronatin | Eutheria | Neural development | (Evans et al. 2005) |
| IGIP | IgA-inducing protein | Eutheria | Activates the production of immunoglobulin A by B cells. | (Endsley et al. 2009) |
| SMCP | sperm mitochondrial-associated cysteine-rich protein | Eutheria | Involved in sperm motility | (Nayernia et al. 2002) |
| CLLU1 | chronic lymphocytic leukemia up-regulated 1 | Primates | Over-expressed in leukemia, *de novo* origin | (Knowles and McLysaght 2009) |
| HMHB1 | histocompatibility (minor) HB-1 | Primates | Precursor of the histocompatibility antigen HB-1, *de novo* origin | (Toll-Riera et al. 2009) |
| DCD | dermcidin | Haplorrhini | Antimicrobial peptide, secreted in the skin | (Schitteck et al. 2001; Toll-Riera et al. 2009) |
| MYEOV | myeloma over-expressed | Haplorrhini | Over-expressed in myeloma, *de novo* origin | (Chen et al. 2015) |
| RP11-429E11.3 | uncharacterized protein | Great apes | *De novo* origin | (Guerzoni and McLysaght 2016) |
| RP11-45H22.3 | uncharacterized protein | Hum/Chimp | *De novo* origin | (Ruiz-Orera et al. 2015) |

### Mammalian-specific genes are enriched in testis

We compared the gene expression patterns in different tissues using human data from GTEx (Ardlie et al. 2015) and mouse data from ENCODE (Pervouchine et al. 2015) (Tables S2-S7). We generated two control gene sets: a set of randomly chosen genes that were not mammalian-specific (’random’) and a collection of genes with homologues in all 34 non-mammalian species used in step 1 of our method (’ancestral’). We found that mammalian-specific genes were strongly enriched in testis. The number of mammalian-specific genes with highest expression in this organ was 50% in human and 40% in mouse, compared to 20% and 13%, respectively, for ‘random’ genes (Fisher test p-value <10^−5^ for both species). The genital fatpad in mouse also showed a significant enrichment of mammalian-specific genes (Fisher test p-value <0.01), although this affected a much smaller percentage of genes (∼ 5%).

We estimated the number of genes that were tissue-specific using a previously described metric (Yanai et al. 2005). The majority of mammalian-specific genes were tissue-specific, whereas this was not the case for the control gene sets (Figure 2A and S3A, for mouse and human; Fisher test p-value < 10^−5^ for pairwise comparisons between mammalian-specific classes and non mammalian-specific classes, in both species). The ‘mam-basal’ (class 2) genes tended to be more tissue-specific than younger gene classes (Fisher test p-value < 0.005 for both human and mouse). Most mammalian-specific genes that were tissue-specific wre preferentially expressed in testis (Figure 2B and S3B).

**Figure 2.**
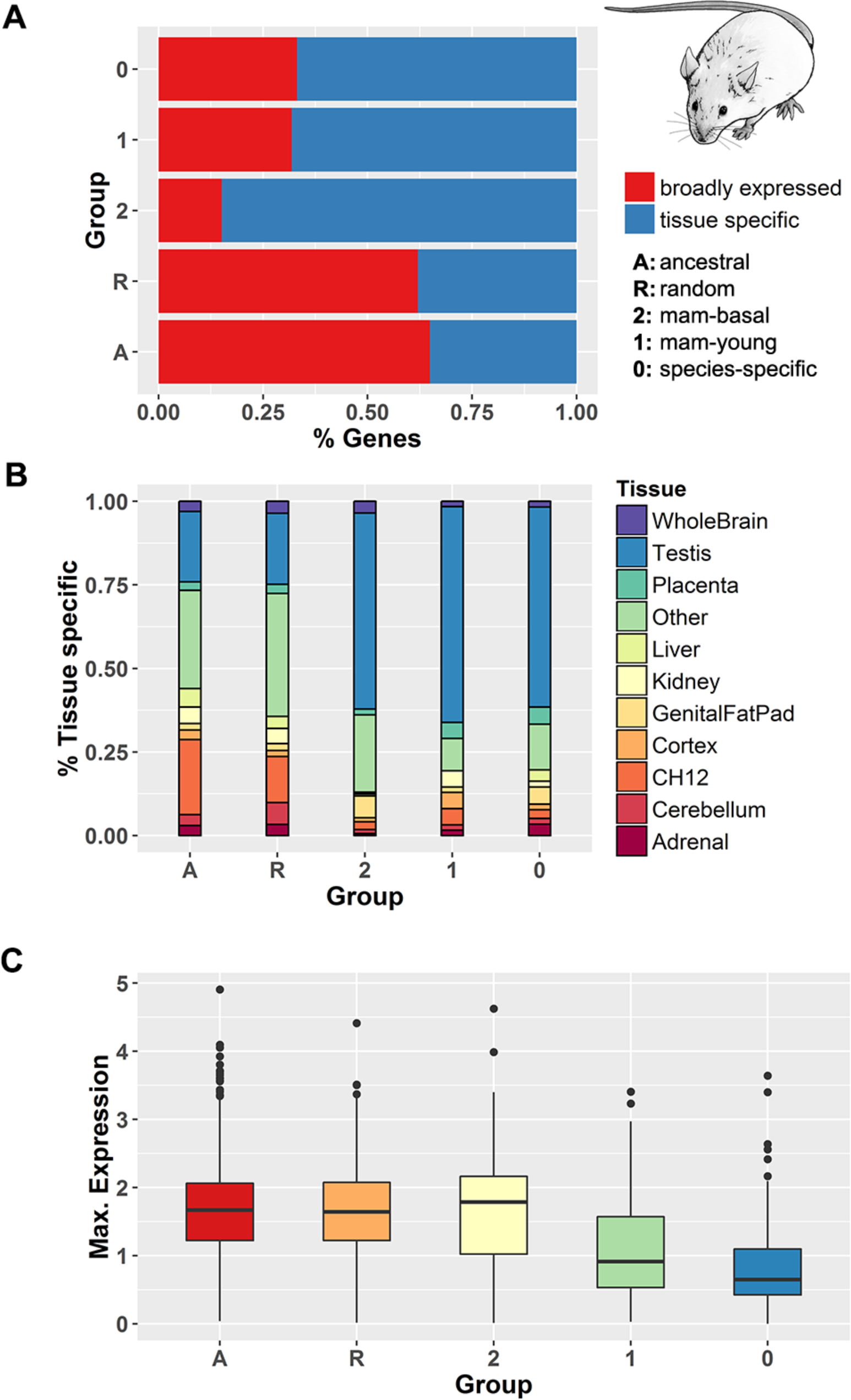
Gene expression patterns of genes from different conservation levels. **A.** Proportion of broadly-expressed and tissue-specific genes in different conservation classes. **B.** Fraction of genes with maximum expression in a given tissue for different conservation classes. **C.** Box-plot showing the distribution of FPKM gene expression values, at a logarithmic scale, in different conservation classes and for the tissue with the highest expression value. Data in B and C is for tissue-specific genes. All data shown is for mouse genes. See Figure S3 for the same data for human genes.

We extracted the gene expression values as FPKMs (Fragments Per Kilobase per Million mapped reads) from GTEx and mouse ENCODE, focusing on the tissue with the highest gene expression. As the three classes of mammalian-specific genes showed similar tissue distributions, their FPKM values could be compared. We found that ‘mam-basal’ genes were, in general, expressed at higher levels than ‘mam-young’ or ‘species-specific’ genes (Wilcoxon test p-value < 10^−4^ for both mouse and human)(Figure 2C and S3C).

An enrichment of mammalian-specific genes in testis was expected given previous observations that nascent transcripts in humans are often expressed in this organ (Ruiz-Orera et al. 2015). However, we expected that the fraction would progressively decrease as we considered older classes. This would fit the ‘out of testis’ hypothesis, which proposes that new genes would be first expressed in testis, probably due to highly permissive transcription in the germ cells (Soumillon et al. 2013), and would later gain expression in other tissues (Vinckenbosch et al. 2006). Contrary to this expectaton, we did not observe any difference between genes presumbaly originated at different times in the mammalian phylogeny. This suggests that not only new genes are born in testis but that many lineage-specific genes may have important functions in this organ.

### Mammalian-specific proteins tend to be short

Different studies have found that proteins encoded by genes with a narrow phylogenetic distribution tend to be shorter than average (Albà and Castresana 2005; Carvunis et al. 2012; Toll-Riera et al. 2012; Neme and Tautz 2013; Arendsee et al. 2014; Palmieri et al. 2014). We investigated whether this trend was also true in our dataset. We compared the mammalian-specific proteins from different conservation levels to the ‘random’ and ‘ancestral’ gene sets. We confirmed that the mammalian-specific proteins were significantly shorter than the other proteins (Figure 3A, Wilcoxon test p-value < 10^−5^). Proteins in the ‘mam-basal’ were slighlty longer than those in the ‘mam-young’ or ‘species-specific’ sets (Wilcoxon test p-value < 0.01). When we restricted the analysis to human or mouse proteins we obtained similar results (Figure S4).

**Figure 3.**
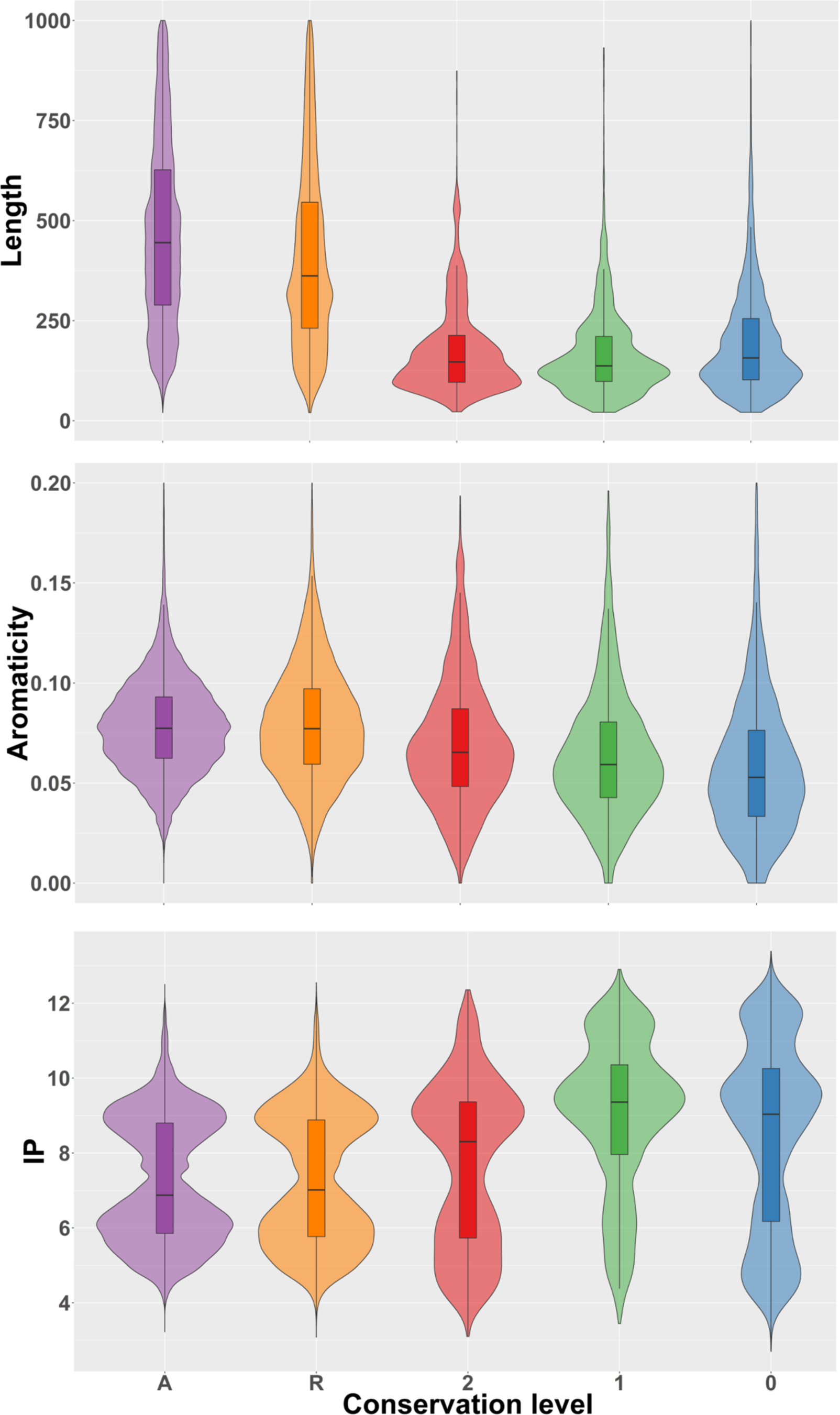
Sequence properties of mammalian-specific genes. **A.** Sequence length in amino acids. **B.** Aromaticity. **C.** Isoelectric point (IP). Protein sequences were extracted from the complete gene families set. We used the following gene groups: A: ancestral; R: random; 2: ‘mam-basal’; 1: ‘mam-young’; 0: species-specific.

### Proteomics support

We used the PRIDE peptide database (Vizcaíno et al. 2016) to search for matches to the sets of human and mouse mammalian-specific proteins. We required at least two unique matching peptides. Using negative controls derived from intronic or random sequences we obtain that these conditions were very stringent and corresponded to only ∼0.2% false discovery rate. Despite the short size of mammalian-specific proteins, potentially hindering their detection by mass spectrometry (Slavoff et al. 2013), we could find proteomics evidence for a large number of them (Figure 4A, Table S8 and S9 for human and mouse, respectively). For example, the number of ‘mam-basal’ mouse proteins with proteomics evidence was 88%, and in the classes ‘mam-young’ and ‘species-specific’ the percentage was also remarkably high (≥ 75%). In most cases this was supported by more than two PRIDE peptides (Figure 4B and 4C). In humans peptide coverage was lower than in mouse, for example about 50% of the human proteins in the ‘mam-basal’ group had peptide hits compared to 88% in mouse. As these proteins should be present in both species, this can be attributed to a higher coverage of mouse peptides with respect to human peptides in PRIDE.

**Figure 4.**
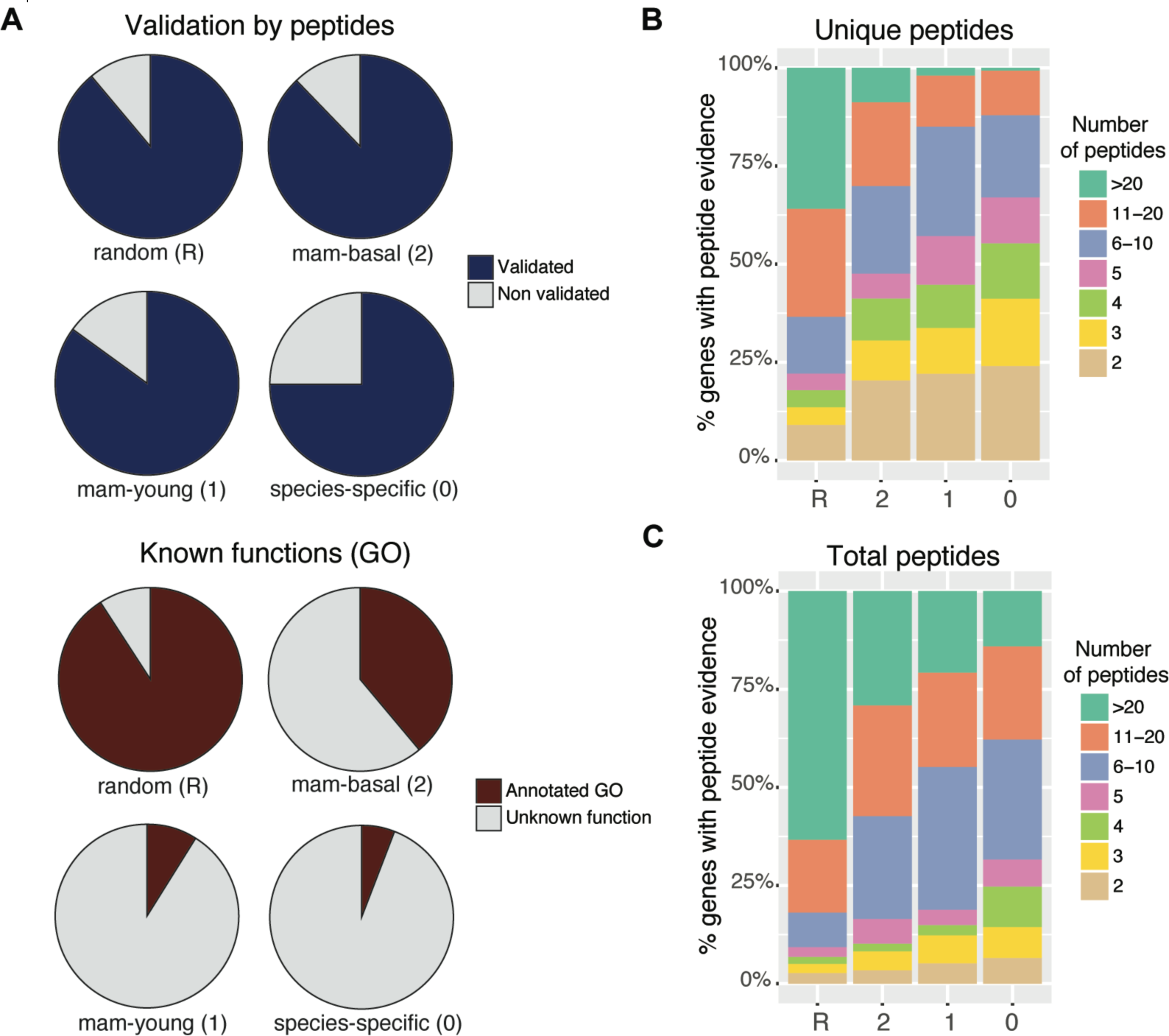
Proteomics and Gene Ontology information. **A.** Proportion of mouse genes with proteomics or Gene Ontology (GO) data for different gene groups. Validated proteins were those that had at least two different peptides with a perfect match and these peptides did not map to any other protein allowing for up to two mismatches. **B.** Number of unique peptides for validated proteins from different groups. **C.** Number of total peptides for validated proteins from different groups.

### Biases in aromaticity and isoelectric point

In addition to sequence length, some studies have reported differences in the sequence composition of young proteins with respect to older proteins (Carvunis et al. 2012; Arendsee et al. 2014; Wilson et al. 2017). Here we inspected the distribution of aromaticity and isoelectric point (IP) values (Figure 3B and 3C, respectively) in the different gene sets. We found that aromaticity values were significantly lower in mammalian-specific proteins than in proteins from the classes ‘ancestral’ and ‘random’ (Wilcoxon test pvalue < 10^−5^). This effect could be clearly appreciated at the different conservation levels. These findings were confirmed in the human and mouse protein subsets (Figure S5).

IP values were significantly higher in mammalian-specific proteins than in ‘ancestral’ and ‘random’ proteins (Wilcoxon test, p-value < 10^−5^). Additionally, among mammalian-specific genes, the most recent families (’mam-young’ and ‘species-specific’) had significantly higher IP values than the oldest ones (’mam-basal’) (Wilcoxon test, p-value < 10^−5^). This indicated a depletion of negatively charged amino acids in the youngest proteins. This trend was also confirmed in the human and mammalian protein subsets (Figure S6).

### Functions of mammalian-specific genes

We searched for information on gene function using Gene Ontology (GO) (Ashburner et al. 2000). We focused on human and mouse because these species are relatively well-annotated and at the same time sufficiently distant as to contain completely different families in the class ‘mam-young’. We found functional data for 38% of the ‘mam-basal’ families, a relatively low percentage when compared to that for more widely conserved genes (’random’ ∼90%)(Tables S10 and S11). But the most striking finding was the low percentage of ‘mam-young’ families with functional data, which was only of about 6% for both human and mouse protein containing families. This anecdotal level of annotation was consistent across the different ‘mam-young’ nodes in the phylogeny. Figure 4A shows the percentage of genes annotated with GO terms for the different mouse conservation groups. There is a striking contrast between the low level of functional annotation and the high level of proteomics support for young mammalian-specific gene families.

We performed a functional enrichment test with DAVID (Huang et al. 2009). Table 2 shows a summary of the main functional groups for human mammalian-specific genes, very similar results were obtained for mouse (Tables S12 and S13). The ‘mam-basal’ class was significantly enriched in terms related to ‘immune response’, ‘reproductive process’ and ‘extracellular region’ (Fisher exact test, Benjamini-Hochberg multiple test correction p-value<0.01). The ‘mam-young’ class was only weakly enriched in the term ‘extracellular region’ (p-value = 0.04). The lack of results in this group was expected given the small percentage of genes with known functions. We also consulted the International Mouse Phenotyping Consortium (IMPC) for targeted knockout data on mouse mammalian-specific genes. We found 11 genes with phenotypes related to pigmentation, abnormal morphology of the seminal vesicle and preweaning lethality, among others (Table S14).

**Table 2.** **Main functions of mammalian-specific genes.** The results shown are for human genes classified as ‘mam-basal’ (class 2).

| Enriched function | Representative terms | N genes | Corrected p-value |
| --- | --- | --- | --- |
| 1. Immune response | 1.1 immune response (GO) | 14 | 2.2E-3 |
|  | 1.2 cytokine activity (GO) | 13 | 1.6E-10 |
|  | 1.3 Jak-STAT signaling pathway (KEGG) | 6 | 5.5E-4 |
| 2. Reproduction | 2.1 reproductive process in a multicellular organism (GO) | 12 | 1.3E-3 |
|  | 2.2 spermatogenesis (GO) | 10 | 9.2E-4 |
| 3. Secreted protein | 3.1 extracellular region (GO) | 64 | 1.8E-15 |
|  | 3.2 secreted (Uniprot) | 59 | 2.8E-14 |
|  | 3.3 signal peptide (Uniprot) | 60 | 7.0E-10 |

We describe the main overrepresented functional classes below.

#### 1. Immune response

Proteins involved in immune response encoded by mammalian-specific genes included several peptides modulating the activity of B or T cells (cytokines) as well as a number of known antimicrobial peptides (AMPs). In the first group there were several interleukins (IL2, IL3, I13, IL32 and IL33), the tumor necrosis factor superfamily 9 and the IgA-inducing protein (IGIP) (see Table 1). AMPs comprised dermcidin (Schittek et al. 2001), mucin-7 (Bobek and Situ 2003) and C10orf99 (Yang et al. 2015).

AMPs are part of the innate immune system. They are small proteins that contain specific sequence patches, often enriched in basic amino acids, that directly interfere with the bacteria or fungus cell membrane. We used the software AMPA (Torrent et al. 2012) to evaluate the AMP sequence propensity of mammalian-specific proteins. This software produces a score that is inversely related to the AMP potential of the protein, and which identifies protein stretches with putative antimicrobial activity. First, we confirmed that human proteins with known AMP activity had lower AMPA scores than proteins for which such activity has not been described (Table S15, Wilcoxon test p-value < 10^−5^). Second, we compared mammalian-specific proteins to a set of proteins of similar size but conserved in other vertebrates (control set, Table S15). We found that the mammalian-specific proteins had significantly lower scores than the control set (Wilcoxon test p-value = 0.0009). We also discovered that the proteins classified as ‘mam-basal’ had an excess of proteins with two or more putative AMP sequence stretches with respect to the control set (Fisher test p-value=0.016). One example was mucin-7, which contained a previously described stretch with anti-fungal activity (Bobek and Situ 2003). The program predicted a second putative sequence with antimicrobial activity that we validated experimentally (Figure S7). Although it is difficult to predict which specific proteins are *bona fide* AMPs based on this data alone, the general enrichment in AMP-like features nevertheless suggests that a number of new proteins may function as AMPs.

#### 2. Reproduction

Genes with the term reproductive process had a varitey of functions related to spermatogenesis or sperm motility. The group included proteins involved in the replacement of histones by protamines in the germ cells, such as transition protein 2. Other proteins had structural roles, such as the sylicins, which are an integral part of the complex cytoskeleton of sperm heads (Hess et al. 1993). Proteins affecting sperm motility included protamine 3 (Grzmil et al. 2008) and the mitochondrion-associated cysteine-rich protein (SMCP) (Nayernia et al. 2002). Mice in which the gene encoding SMCP has been disrupted exhibited reduced motility of the spermatozoa and decreased capability of the spermatozoa to penetrate oocytes (Nayernia et al. 2002). In general, many of the mammalian-specific genes showed the highest level of expression in testis, suggesting that a substantial fraction of them is likely to have reproduction-related functions.

#### 3. Secreted proteins

A large group of mammalian-specific gene families were annotated as “secreted protein” and/or “extracellular space” (Table 2). This group included many proteins from the immune response system but also proteins secreted in other organs, such as the mammary gland (caseins), skin (dermokine) or the lung (secretoglobins). This highlights the importance of secreted or extracellular molecules in recent mammalian adaptations.

### Origin of mammalian-specific proteins

Mammalian-specific protein-coding genes may derive from already existing coding sequences or may have originated *de novo* from previously non-coding genomic regions (Toll-Riera et al. 2009; Neme and Tautz 2013). In the context of new genes for which ancestors cannot be traced back using standard sequence similarity searches, as described in this paper, the first process often involves complex sequence rearrangements and the co-option of genomic DNA segments as new coding exons. Comparative genomics studies have shown that the caseins, which transport calcium in milk, probably originated from genes encoding teeth proteins during the early evolution of mammals, following a series of gene duplications, sequence rearrangements and rapid sequence divergence events (Kawasaki et al. 2011). Another example is the mammalian-specific Late Cornified Envelope (LCE) group of proteins. The LCEs are part of a gene cluster shared by mammals, birds and reptiles, known as the epidermal differentiation complex (EDC). In this cluster, multiple episodes of sequence duplication and divergence have resulted in the extraordinary functional diversification of epidermal proteins in mammals (Strasser et al. 2014).

In other cases the genes may have originated *de novo* from a previously non-coding sequence of the genome (Begun et al. 2006; Levine et al. 2006; Toll-Riera et al. 2009; Knowles and McLysaght 2009; Wu et al. 2011; Murphy and McLysaght 2012; Samusik et al. 2013; Chen et al. 2015; Ruiz-Orera et al. 2015; Guerzoni and McLysaght 2016). The definition of a *de novo* gene usually includes the requirement that the syntenic genomic regions from closely related species lack a coding sequence of a similar length. This means that *de novo* genes that originated a long time ago will be very difficult to identify, as genome synteny information will no longer be available. Our dataset contained 15 human and 13 mouse *de novo* protein-coding genes identified in previous studies (Tables S16 and S17, respectively; Table 1 for selected examples). One example was the human minor histocompatibility protein HB-1, which modulates T-cell responses and previously defined as primate-specific (Toll-Riera et al. 2009). As expected, these genes corresponded to recent nodes, containing one or a few closely related species.

### Discussion

The sequencing of complete genomes has resulted in a more accurate view of the number of genes present in each species and how these genes relate to genes in other species. A puzzling discovery has been that a sizable fraction of genes does not have homologues in other species (Toll-Riera et al. 2009; Donoghue et al. 2011; Tautz and Domazet-Lošo 2011; Macarena Toll-Riera et al. 2011; Carvunis et al. 2012; Wissler et al. 2013). A well-known mechanism for the formation of new genes is gene duplication (Ohno 1970). However, the evolution of gene duplicates is limited by the structural and functional constraints inherited from the parental gene (Ohno and Epplen 1983; Pegueroles et al. 2013; Pich I Roselló and Kondrashov 2014). Sequences that were not previously coding but that have been coopted for a coding function, called *de novo* genes, are free from such constraints (Levine et al. 2006; Knowles and McLysaght 2009; Xie et al. 2012; Ruiz-Orera et al. 2014). Genes with new coding sequences are likely to drive important species- and lineage-specific adaptations (Khalturin et al. 2009; Johnson and Tsutsui 2011) but we still know very little about their specific functions.

In order to advance our knowledge in this field we have generated a comprehensive set of mammalian-specific gene families and analysed the properties of genes conserved at different levels in the mammalian phylogeny. For this we have computed extensive sequence similarity searches using both annotated proteomes and RNA-Seq-derived data, and each family has been assigned to one of the 29 internal nodes or 30 terminal branches of the tree. This is different from the classical “phylostratigraphy” approach, which is based on the gene homologues that we can detect for a given species in a set of other species (Albà and Castresana 2005; Domazet-Lošo et al. 2007). The integration of data from multiple species is expected to result in a more robust classification of the node of origin of each gene, and the gene families that are specific to each group of organisms can be directly retrieved and studied. One important limitation of this and other studies is the lack of direct protein quantification data for most species. Therefore, the gene families were mostly based on coding sequence predictions, as extracted from the gene annotations, as well as transcriptomics data.

In mammals, the skin needs to be flexible and thin enough to allow muscles to produce an elastic deformation. The outermost layer of the skin, known as *stratum corneum*, is particularly flexible in this group when compared to the stiff skin of reptiles (Alibardi 2003). Several mammalian-specific genes detected in this study were involved in the formation of skin structures, such as the Late Cornified Envelope (LCE) family, Dermokine, Keratinocyte Differentiation-Associated Protein, and Corneodesmosin (CDSN). The latter protein participates in the specialized junctions known as corneodesmosomes, which bridge together corneocytes in the lower part of the *stratum corneum* (Jonca et al. 2011). Additionally, we found a gene associated with psoriasis (PSOR-1), localized in the same genomic region than CDSN. Another important adaptation in mammals is the production of milk; in most mammals, the most abundant milk proteins are caseins, which form micelles that transport calcium. Our method identified several caseins (alpha S1, beta, kappa), which are part of a larger family of secretory calcium-binding phosphoproteins (SCPP) (Kawasaki et al. 2011).

Another large group of mammalian-specific genes encoded proteins involved in the immune response. One example was the IgA-inducing protein (IGIP), a short secreted protein which is composed of only 52 residues. This protein is produced in the dendritic cells and stimulates the production of immunoglobulin A in B cells (Endsley et al. 2009). Although its sequence is highly conserved across mammals, no homologues have been found in other vertebrates. Our set of mammalian-specific genes also included several previously described antimicrobial peptides (AMPs). The antimicrobial active regions in AMPs are often enriched in certain residues, including arginines and cysteines (Yeaman and Yount 2007). This can be used to predict the propensity of a protein to be an AMP (Torrent et al. 2012). We observed higher AMP propensities in mammalian-specific genes than in non mammalian-specific genes, suggesting that there may be additional mammalian-specific AMPs that have not yet been characterized.

A study that used EST data to determine tissue gene expression patterns found that retrogenes were enriched in testis when compared to multiexonic genes (Vinckenbosch et al. 2006). The frequent expression of young genes in testis led to the ‘out of testis’ hypothesis, which proposes that new genes would be initially expressed in testis and would later evolve broader expression patterns (Kaessmann 2010). A subsequent study that used high throughput RNA sequencing data from several mammalian species and tissues found that the proportion of testis-specific retrogenes decreased with gene age, providing further support to this hypothesis (Carelli et al. 2016). In our study, however, we found a similar proportion of testis-specific genes for different mammalian-specific age classes. This enrichment probably reflects the importance of sexual selection in driving changes in the reproductive organs, both at the anatomical and molecular levels (Gage et al. 2004; Kleene 2005). New genes that increase sperm competitiveness will rapidly reach fixation in the population, and subsequently be preserved by purifying selection.

A number of gene properties, including gene expression tissue specificitiy and protein length, have been previously shown to correlate with gene conservation level (Albà and Castresana 2005; Carvunis et al. 2012; Zhang et al., 2012; Neme and Tautz, 2013). It is easy to understand why new coding sequences, specially those originated *de novo*, will tend to be shorter than proteins that have been evolving for a long time. Open reading frame (ORF) generated by chance are often shorter than 100 amino acids. In contrast, proteins which have evolved over extended time periods usually contain several proteins domains, which may have been gained at diferent times (Buljan et al. 2010; Toll-Riera and Albà 2013; Andreatta et al. 2015). We also identified a number of trends that did not show a linear correlation with gene age. For example old genes were more broadly expressed than mammalian-specific genes, but ‘mam-young’ or ‘species-specific’ were also more broadly expressed than ‘mam-basal’ genes. We also observed that mammalian-specific proteins, specially those in the two youngest classes, were significantly depleted of negatively charged residues. The reasons for this bias remain enigmatic but we speculate that it may be related, at least partly, to some of these proteins having a yet uncharacterized antimicrobial activity.

BLASTP has been shown to be very sensitive to detect homologues between closely related species when the proteins are evolving at a constant rate (Albà and Castresana 2007). However, very drastic sequence changes can occur during functional shifts, compromising the capacity of BLAST to detect homologues. As already mentioned, we found several cases which corresponded to a model of gene duplication followed by very rapid sequence divergence and neofunctionalization (Kawasaki et al. 2011; Grayson and Civetta 2012; Strasser et al. 2014). Other cases were *de novo* genes previously reported in the literature. The vast majority of the youngest genes had unknown functions but were nevertheless supported by proteomics data. We identified 83 human- or primate-specific genes with proteomics evidence. This number was based on peptides stored in the PRIDE database and was more than one order of magnitude higher than that previously obtained by Ezkurdia and co-workers using other sources of data (Ezkurdia et al. 2014). The controls we performed indicated that our pipeline was highly specific and thus it seems very likely that the proteins are actually produced.

Species-specific genes were surprisingly abundant in comparison to other classes. The number was variable depending on the species, which is expected given that the genomes have been annotated by different research consortia. For example, whereas for most primate genomes the annotations are primarily based on the human genome, in other species, such as the lynx, extensive RNA-Seq data has also been employed (Abascal et al. 2016). Additionally, the rate of molecular evolution varies considerably in different groups. For instance the distance between mouse and rat is of about 0.2 substitutions/site, which is comparable to the distance separating the most divergent primate species. It is thus not surprising that mouse had a very larger number of species-specific genes than human; we found proteomics evidence for 291 of the mouse-specific genes. An excess of species-specific genes in phylostratigraphy-based studies has been previously observed (Neme and Tautz 2013). These observations are consistent with a high rate of *de novo* gene emergence accompanied by frequent gene loss in the first stages of the evolutionary history of a gene (Neme and Tautz 2014; Palmieri et al. 2014). We also have to consider that some of the very young genes may not be functional even if translated (Ruiz-Orera et al. 2016).

Contrary to early predictions (Casari et al. 1996), the sequencing of new genomes has not solved the mystery of orphan genes (genes for which we find no homologues in other species); in fact, we now have an ever larger number of orphan genes than we did before. The cell expresses many transcripts with low coding potential, or long non-coding RNAs, which are species- or lineage-specific, and which can potentially be translated (Ruiz-Orera et al. 2014). Whereas there is ample evidence for continuous new gene emergence, it is unclear which functions new genes eventually contribute to. On the basis of over 400 mammalian-specific genes that are relatively well-conserved and have functional annotations we have found that many new genes encode secreted proteins and that their formation may have been advantageous in the response against pathogens or mating. The collection of gene families obtained here will help accelerate future studies on the evolutionary dynamics and functions of novel genes.

## MATERIALS AND METHODS

### Sequence sources

We initially identified 68 mammalian species that had fully sequenced genomes and which showed a relatively sparse distribution in the mammalian tree. The proteomes and cDNA sequences were downloaded from Ensembl v.75 (Flicek et al. 2014), the National Center for Biotechnology Institute (NCBI Genbank version available on March 2014)(Benson et al. 2015) and the *Lynx pardinus* Genome Sequencing Consortium (Abascal et al. 2016). The tree topology, and the approximate number of substitutions per site in each branch, were obtained from previous studies (Meredith et al. 2011; Toll-Riera et al. 2011; O’Leary et al. 2013). We downloaded RNA sequencing (RNA-Seq) data for 30 different species using the public resource Gene Expression Omnibus (GEO) (Barrett et al. 2013). The RNA-Seq samples were from different body tissues depending on the species. The total number of RNA-Seq samples was 434, with a median of 10 samples per species.

### Identification of mammalian-specific genes

We run BLASTP sequence similarity searches for each of the mammalian proteomes against a set of 34 non-mammalian proteomes (Table S1). All BLASTP searches were run with version 2.2.28+ using an evalue threshold of 10^−4^ and the filter for low complexity regions (LCRs) activated (seg = ‘yes’) (Altschul et al. 1997). Consequently, we did not consider proteins that were extensively covered by LCRs (with less than 20 contiguous amino acids devoid of LCRs as measured by SEG with default parameters (Wootton and Federhen 1996)). We also discarded genes encoding a protein that had significant sequence similarity to non-mammalian species, as well as genes that had paralogues with homologues in non-mammalian species, as previously described (Toll-Riera et al. 2009).

We generated two control sets of non mammalian-specific genes. The first set, which we named ‘ancestral’ (A) contained proteins that had homologues in all the 34 non-mammalian species mentioned above. This set of genes is well-conserved across eukaryotes, having originated in a common ancestor. The second set, named ‘random’ (R), contained 1,000 proteins (for each species) that were not in the mammalian-specific group.

### Building gene families based on gene annotations

We built a mammalian tree for the species with complete genomes using previous published data (Meredith et al. 2011; O’Leary et al. 2013). The next step was to develop a pipeline to construct gene families and to assign them to a node in the tree. We wanted the gene families to be as large as possible and include both orthologues and paralogues. The node represented the point from which no further ancestors could be traced back.

We first ran BLASTP searches for every set of mammalian-specific genes in each species against the mammalian-specific genes in the other species (e-value<10^−4^). This resulted in a node of origin for each individual gene and a list of homologous proteins in the same and other species. Next we collated the information for all the homologous proteins, progressively expanding the gene families until no more members could be added (single linkage clustering). The group then was assigned to the oldest possible node considering the species present in the gene family.

### Transcript reconstructions from RNA-Seq data

The RNA-Seq sequencing reads of each sample underwent quality filtering using ConDeTri (v.2.2) (Smeds and Künstner 2011) with the following settings (−hq=30 −lq=10). Adaptors were trimmed from filtered reads if at least 5 nucleotides of the adaptor sequence matched the end of each read. In all paired-end experiments, reads below 50 nucleotides or with a single pair were not considered. We aligned the reads to the corresponding reference species genome using Tophat (v. 2.0.8) (Kim et al. 2013) with parameters −N 3, −a 5 and –m 1, and including the corresponding parameters for paired-end and stranded reads if necessary. We performed gene and transcript assembly with Cufflinks (v 2.2.0) (Trapnell et al. 2010) for each individual tissue. We only considered assembled transcripts that met the following requirements: a) the transcript was covered by at least 4 reads. b) transcript abundance was >1% of the abundance of the most highly expressed gene isoform. c) <20% of reads were mapped to multiple locations in the genome. d) The reconstructed transcripts were at least 300 nucleotides long. Subsequently, we used Cuffmerge to build a single set of assembled transcripts for each species. We use Cuffcompare to compare the coordinates of our set of assembled transcripts to gene annotation files from Ensembl (gtf format, v.75) or NCBI (gff format, December 2014), to identify annotated transcripts and, to generate a set of novel, non-annotated, transcripts.

### Refinement of the age of the families using transcriptomics data

We performed TBLASTN (e-value < 10^−4^) searches of the proteins in each gene family against the set of novel transcripts assembled from RNA-Seq samples. If we found any homologous sequence in a species that was more distant to the members of the family than the originally defined node we reassigned the family accordingly, to an earlier node. This was based on conservative criteria, the presence of an homologous expressed sequence, even if not annotated as a gene, was considered as evidence that the gene had originated at an earlier node connecting all homologous transcripts.

In order to minimize the biases due to the different availability of RNA-Seq data in different parts of the tree we decided to focus only on the 30 species with RNA-Seq data for all analyses. Besides, we only considered families with sequence representatives in at least half of the species derived from the node to which the family had been assigned. The use of transcriptomics data filled many gaps in the tree and resulted in a deepening of the node of origin in some cases. The use of RNA-Seq data allowed us to expand the range of species for 1,126 of the gene families initially defined as species-specific (∼22%) and to increase the number of species per family in 617 multi-species families (68%). The procedure resulted in 2,013 multi-species gene families and 3,972 species-specific gene families (mostly singletons).

### Gene expression data

We retrieved gene expression data from GTEx (Ardlie et al. 2015) and mouse ENCODE (Stamatoyannopoulos et al. 2012) tissue expression panels. We only analyzed genes which were expressed in at least one tissue sample. We computed the FPKM (Fragments per Kilobase Per Million mapped reads) median expression for each gene and tissue available in GTEX (release 2014, Ensembl v.74). For Mouse ENCODE, which was based on Ensembl v.65 (mm9), we computed the FPKM mean for each gene and for tissues with 2 replicas we only used the data if reproducibility indexes were lower than 0.1. We identified the tissue with the highest expression value and calculated the tissue preferential expression index as previously described (Yanai et al. 2005; Ruiz-Orera et al. 2015). Genes for which this index was 0.85 or larger were classified as tissue-specific.

### Sequence features and proteomics data

The analysis of the sequences from different sets was performed using Python scripts. We employed Biopython embedded functions to calculate the isoelectric point (IP) and aromaticity. We used the proteomics database PRIDE (Vizcaíno et al. 2016) to search for peptide matches in the proteins encoded by various gene sets. For a protein to have proteomics evidence we required that it had at least two distinct peptide perfect matches and that the peptides did not map to any other protein allowing for up to two mismatches. We estimated the false discovery rate was below 0.2% using two different negative control sets. The first one consisted in building fake sets of human and mouse protein sequences, by preserving the amino acid composition and protein length of the ‘random’ human and mouse datasets, and subsequently searching for peptide matches. After repeating the procedure 10 times, we obtained that 0.18 % of the human proteins, and 0.69% of the mouse proteins had 1 or more peptide hits, but none of them had 2 or more peptide hits. The second control used translated intronic sequences of the same length as the coding sequences in the ‘random’ human and mouse datasets. After repeating the procedure 10 times, we obtained that 0.36 % of the human proteins, and 1.68 % of the mouse proteins had 1 or more peptide hits, but only 0% and 0.19% of the human and mouse proteins, respectively, had 2 or more peptide hits.

### Functional analysis

We downloaded Gene Ontology (GO) terms from Ensembl v.75 for all human and mouse genes (Tables S11 and S12, respectively). In order to estimate the number of gene with known functions in each conservation class we counted how many genes were associated with at least one GO term. In this analysis we did not consider the terms ‘biological process’, ‘nucleus’, ‘cellular_component’, ‘molecular_function’, ‘cytoplasm’, ‘plasma membrane’, ‘extracellular space’, ‘protein binding’, ‘extracellular region’ or ‘integral to membrane’, as they were very general and did not imply knowledge of the specific function of the protein.

We used DAVID (Huang et al. 2009) to assess enrichment of particular functions or subcellular locations in mammalian-specific genes from human and mouse (Tables S13 and S14, respectively).

### Prediction of antimicrobial activity

We wanted to test if mammalian-specific genes were enriched in AMP-like features. This was motivated by the enrichment of immune response proteins among mammalian-specific genes (Table 2), including three known AMPs (dermcidin, mucin 7, C10orf99 protein) and the skew towards high isoelectric point values observed in these proteins (Figure 3). We measured the antimicrobial activity potential of all human proteins using the program AMPA (Torrent et al. 2012). In the majority of proteins from a set of 59 genes encoding known AMPs, which we gathered from the literature, AMPA was able to predict stretches with AMP activity (Table S15, AMP status ‘known’). We then used this program to calculate scores and number of putative AMP stretches in mammalian-specific proteins (Table S15, type ‘mammalian-specific’) as well as in a large set of size-matched non mammalian-specific proteins (Table S15, type ‘conserved’).

The peptides for testing the antimicrobial activity were prepared by Fmoc solid phase synthesis methods, purified by HPLC and characterized by mass spectrometry, as previously described (Falcao et al. 2015). We assayed the activity of mucin-7 peptideson reference strains of *E. coli* (ATCC 25922), *P. aeruginosa* (ATCC 27853), *E. faecalis* (ATCC 29212), and *S. aureus* (ATCC 29213) at the Microbiology Service of Hospital Clínic (Universitat de Barcelona). Minimal inhibitory concentration (MIC) assays were performed by the microdilution method in Mueller–Hinton broth according to Clinical and Laboratory Standards Institute (CLSI) guidelines. As a positive control the Cecropin A-Melittin peptide CA(1-8)M(1-18) was employed (Saugar et al. 2002). This peptide exhibited MIC values ranging from 0.5 to 64 in the four bacteria strains tested.

### Statistical Data Analyses

We used Python 2.7 to parse the data from different programs and files, cluster the genes into gene families, and calculate sequence-based statistics. The ete2 package was used to perform analyses using the phylogenetic tree structure (Huerta-Cepas et al. 2010). We generated plots and performed statistical tests with R (R Development Core Team 2013).

## SUPPLEMENTARY MATERIAL

Supplementary figures can be found in Supplementary file 1. Supplementary tables can be accessed at https://figshare.com/articles/Villanueva-Ca_as_etal_supTables/4542949. The list of mammalian-specific gene families is available at http://dx.doi.org/10.6084/m9.figshare.4239404. Protein sequences for the gene families are available at http://dx.doi.org/10.6084/m9.figshare.4239440. Information on the RNASeq datasets is available at http://dx.doi.org/10.6084/m9.figshare.4239431. Transcripts assemblies generated from the RNA-Seq data and used in node of origin estimation are available at http://dx.doi.org/10.6084/m9.figshare.4239449.

## ACKNOWLEDGEMENTS

We acknowledge Sarah Djebali for providing us with data on mouse Encode, Roger Hall and Sara Díez for the beautiful drawings in Figures 1 (RH) and 2 (SD), Javier Valle for help with peptide synthesis and William Blevins for useful comments on the manuscript. The work was funded by grants BFU2012-36820 and BFU2015-65235-P from Ministerio de Economía e Innovación (Spanish Government) and co-funded by FEDER. We also received funding from Agència de Gestió d’Ajuts Universitaris i de Recerca Generatlitat de Catalunya (AGAUR), grant number 2014SGR1121.

